# A transgenic pig model expressing a ZsGreen1 reporter across an extensive array of tissues

**DOI:** 10.1101/293258

**Authors:** Amy T. Desaulniers, Rebecca A. Cederberg, Elizabeth P. Carreiro, Channabasavaiah B. Gurumurthy, Brett R. White

## Abstract

**Background:** The advent of genetically engineered pig production has revealed a wide array of opportunities to enhance both biomedical and agricultural industries. One powerful method to develop these models is transgenesis; however, selection of a suitable promoter to drive transgene expression is critical. The cytomegalovirus (CMV) promoter is the most commonly used viral promoter as it robustly drives transgene expression in a ubiquitous nature. However, recent reports suggest that the level of CMV promoter activity is tissue-dependent in the pig. Therefore, the objective of this study was to quantify the activity of the CMV promoter in a wide range of porcine tissues. Swine harboring a CMV-ZsGreen1 transgene with a single integration site were utilized for this study. Thirty five tissue samples were collected from neonatal hemizygous (n = 3) and homozygous (n = 3) transgenic piglets and analyzed for ZsGreen1 abundance via immunoblot.

**Results:** ZsGreen1 was detected in all tissues examined; however, quantification revealed that ZsGreen1 protein levels were tissue-specific. Within organs of the digestive system, for example, ZsGreen1 was most abundant in the salivary gland, moderately produced in the esophagus and levels were lowest in the stomach. Interestingly, abundance of ZsGreen1 also differed within organ. For instance, levels were highest in the right ventricle compared with other chambers of the heart. There was no effect of transgene dose as ZsGreen1 expression patterns were similar between homozygous and hemizygous piglets.

**Conclusions:** Ultimately, these results elucidate the tissue-specific activity of the CMV promoter in the neonatal pig. Moreover, this model can serve as a useful tool for research applications requiring reporter gene activity in mammalian organs.

## Background

Use of genetically engineered pig models has grown dramatically in recent years [1]. Transgenic swine are increasingly being utilized as animal models for human disease since they are anatomically, physiologically and phylogenetically more similar to humans than rodents [1, 2]. Consequently, successful xenotransplantation of transgenic pig organs to humans is eminent [3]. In addition to biomedical applications, genetically engineered pigs are also being developed to better understand porcine physiology and enhance the agricultural industry [4-7].

The first genetically engineered pig was generated in an effort to enhance growth rates and harbored a transgene containing the mouse metallothionein-I promoter fused to the human growth hormone gene [8]. Since then, new methodologies (e.g., homologous recombination, gene editing) have enhanced the ease and efficiency of modifying the pig genome [9]. However, incorporation of a transgene remains a prominent method to produce genetically engineered pigs [1]. The most critical aspect of transgenic animal production is promoter selection [10]. Depending on the research interest and application, promoter types include tissue-specific, cell-specific, inducible/conditional or ubiquitous [10]. Since ubiquitous promoters drive transgene expression in all tissues, they are a popular choice to address whole-body questions [10].

The human cytomegalovirus (CMV) immediate early promoter is one example of a ubiquitous promoter. It was discovered by Boshart et al. [11] and has since been heavily utilized to drive recombinant gene expression due to its constitutive and promiscuous nature [12]. These characteristics are directly related to its role in viral infection. Cytomegalovirus is a human herpesvirus which typically lays dormant in infected host cells [13]. Cellular infection quickly (< 1 h) leads to CMV gene expression via the interaction of host transcription factors with the CMV promoter [13]. Gene products of this initial interaction mediate the activation of other promoters within the CMV genome that facilitate virus replication and invasion [13]. Therefore, the CMV promoter has evolved to contain binding sites for many ubiquitous transcription factors in order to replicate in a broad range of host cells [12]. This discovery has had a major impact on the scientific community; the CMV promoter is now the most commonly used ubiquitous promoter and it is also considered the most robust [14].

The CMV promoter has been utilized to drive transgene expression in numerous vertebrate animal models including: sheep [15-17], fish [14, 18], mice [19-22], rats [23, 24], chickens [25] and pigs [4, 26-28]. In the pig, the CMV promoter was used to drive expression of green fluorescent protein (GFP) [27]. As anticipated, expression of the transgene was detected in a wide range of organs, including those derived from each of the 3 germ layers: skin (ectoderm), pancreas (endoderm), and kidney (mesoderm). Despite apparent universal transgene expression [27], the CMV promoter was preferentially active in exocrine vs. non-exocrine cells of the pig [26]. Given the growing interest in the development of transgenic swine models, the objective of this study was to quantify activity of the CMV promoter in a wide range of porcine tissues.

## Methods

### Animals

All animal procedures were conducted in accordance with the University of Nebraska-Lincoln (UNL) Institutional Animal Care and Use Committee. Experiments were performed in the UNL Animal Science Building (Lincoln, NE) and utilized transgenic swine that express ZsGreen1 controlled by the human CMV immediate early promoter; these animals were generated as previously described [4]. In hemizygous animals, one copy of the CMV-ZsGreen1 transgene is stably integrated on chromosome 14 [4], and has remained transcriptionally active throughout the lifetime of the founder animal and in 10 subsequent generations.

Adult animals were housed individually with *ad libitum* access to water and fed approximately 2.5 kg of feed daily. Hemizygous CMV-ZsGreen1 transgenic sows were artificially inseminated with semen from non-littermate hemizygous transgenic boars and allowed to gestate to term. Following Mendelian inheritance, these matings yielded hemizygous, homozygous (2 copies of the transgene), and non-transgenic (control) littermates. After farrowing, piglets remained with their dam to suckle *ad libitum*.

At 1 d of age, 3 littermate female piglets (control, hemizygous transgenic and homozygous transgenic) were selected from 3 different litters; each litter was produced from a different dam and sire mating to maximize genetic diversity. Initially, transgene status was assessed via evaluation of ZsGreen1 expression using an ultraviolet (UV) light and a Roscolux #15 filter (deep straw; Rosco, Port Chester, NY).

### Tissue collection

Piglets were euthanized via intracardiac injection with Euthasol (1 mL/4.53 kg BW; Delmarva Laboratories, Midlothian, VA) and stored at 4°C overnight prior to dissection. The following morning, 35 organ samples generally representing 5 anatomical regions (brain, thoracic, digestive, renal and reproductive) were collected from each transgenic piglet. Isolated tissues included: cerebrum, cerebellum, spinal cord, anterior pituitary gland, posterior pituitary gland, hypothalamus, thymus, lymph node, the four chambers of the heart, lung, muscle, thyroid, salivary gland, esophagus, stomach, duodenum, jejunum, ileum, colon, gall bladder, spleen, pancreas, liver, kidney (medulla and cortex), bladder, adrenal gland (medulla and cortex), ovary, uterus, oviduct, and skin. Skin samples were collected from control littermate piglets to serve as a negative control for ZsGreen1 immunoblotting. In addition, tail samples were collected from all piglets for genotyping. All samples were frozen and stored at −80°C until analysis.

### Deoxyribonucleic acid extraction

Genomic deoxyribonucleic acid (DNA) was isolated from tail samples using an enzymatic digestion and phenol/chloroform extraction. Briefly, a tail slice (approximately 1 mm) was incubated overnight in a lysing solution [50 mM Tris pH 8.0, 150 mM NaCl, 10 mM EDTA, 1% sodium dodecyl sulfate (SDS), 0.1 mg/ml Proteinase K (Fisher Bioreagents, Fairlawn, NJ), 1% β-mercaptoethanol] at 55°C. The samples were extracted twice with a 25:24:1 phenol:chloroform:isoamyl alcohol solution and once with a 24:1 chloroform:isoamyl alcohol solution before precipitating the DNA with isopropanol. DNA was subsequently resuspended in nuclease-free water.

### Genotyping

Since the transgene integration site was previously identified in this model [4], copy number was evaluated via conventional polymerase chain reaction (PCR) using 3 primers. The forward (F) and reverse (R) primers were designed to flank the insertion site of the transgene (T) and an additional reverse primer (RT) aligning the transgene was included (F - GCA ACC TCT TCG ACA CTC CA; R - AGC TAC CAG GGA ACA AAG CC; RT - GGT TTC CCT TTC GCT TTC AAG T). An MJ Research PTC-200 Thermocycler (Waltham, MA) was used for PCR reactions with the following conditions: 1 X *Taq* Buffer A (Fisher Bioreagents) with a final concentration of 2.0 mM MgCl_2_, 200 µM dNTPs, 300 nM of each primer and 1.25 U of *Taq* DNA Polymerase (Fisher Bioreagents). Cycling conditions were 95°C for 2 min, followed by 35 cycles of 95°C for 30 sec, 56°C for 30 sec and 72°C for 60 sec and a final extension of 72°C for 10 min. The resultant PCR products were subjected to electrophoresis on a 1% agarose gel. Products generated from the F and R primer pair were 627 bp in length and reflected a wild type chromosome 14. Products from the F and RT primer pair resulted in an 847 bp product, and indicating that the transgene was present.

### Protein preparation

Protein extraction and sample preparation was performed as described previously [29]. Protein was extracted by homogenization of tissue samples in radioimmunoprecipitation assay (RIPA) buffer (1 ml/100 mg; 20 mM Tris, 137 mM NaCl, 10% glycerol, 1% NP40, 0.1% SDS, 0.5% deoxycholic acid, 2 mM EDTA, 1mM PMSF, 1% protease inhibitor cocktail and 1% phosphatase inhibitor cocktail) using a Biospec Tissue Tearor (Bartlesville, OK). Protein concentrations were quantified using a bicinchoninic acid (BCA) assay (Pierce Biotechnology, Rockford, IL) according to the manufacturer’s instructions. Extracted protein was mixed with 4X loading dye [2% Tris (pH 6.8), 28% glycerol, 20% SDS and Orange G] containing dithiothreitol (DTT; 100 mM) and frozen at −80°C until immunoblotting.

### Immunoblotting

Immunoblotting was performed as described previously [29]. Tissue samples were segregated by anatomical region (brain, thoracic, digestive, renal and reproductive) and protein (10 or 20 µg) was separated using SDS-PAGE (15%) before transfer to an Immobilon-FL polyvinylidene difluoride (PVDF) membrane (Millipore, Billerica, MA) pre-soaked in methanol. After electrophoretic transfer, nonspecific binding was blocked by incubation of blots in Odyssey Blocking Buffer (cat #: 927-40100; LI-COR Biosciences, Lincoln, NE) for approximately 3 h at room temperature. Membranes were then incubated with a mouse monoclonal primary antibody directed against ZsGreen1 (1:1000; cat #: 632598; Clontech, Mountain View, CA), diluted in Odyssey Blocking Buffer with 0.05% Tween-20 and shaking at 4°C overnight. The following morning, membranes were rinsed in Tris-buffered saline with 0.05% Tween-20 (TBS-T) prior to incubation with a goat anti-mouse secondary antibody (1:15000; IRDye 680; #926-32220; LI-COR Biosciences) in Odyssey Blocking Buffer plus 0.05% Tween-20 and 0.025% SDS for 1 h at room temperature. Blots were rinsed with TBS-T to remove excess secondary antibody and briefly rinsed in TBS to remove Tween-20 prior to imaging.

### Quantification of ZsGreen1 abundance

Blots were scanned on an Odyssey Infrared Imager (700 channel; intensity 5.0; 169 micron resolution; LI-COR Biosciences) and converted to greyscale for analysis. Band density was quantitated with Odyssey imaging software (version 2.1; LI-COR Biosciences). Since the abundance of common loading controls (e.g., β-actin) differs across tissue types, total protein levels within each lane were used as the loading control according to Eaton et al. [30]. After ZsGreen1 imaging, membranes were stained with GelCode Blue Safe Protein Stain (ThermoScientific, Rockford, IL) for approximately 1 h and then rinsed with a 50% methanol and 1% acetic acid solution for 1 h. Blots were reimaged (700 channel; intensity 1.0; 169 micron resolution; LI-COR Biosciences), converted to greyscale and quantitated using Odyssey imaging software (version 2.1; LI-COR Biosciences) to determine the relative quantity of protein present in each sample. Data are presented as the relative band density of ZsGreen1 normalized by the total protein levels within the entire lane as described previously [30].

### Statistical Analysis

Analyses were performed using the Statistical Analysis System (SAS; Cary, NC; version 9.4). Analysis of variance (ANOVA) was performed via the MIXED procedure of SAS with tissue type as the fixed effect. Blot and litter were random effects. *P* < 0.05 was considered significant, whereas *P* < 0.10 was considered a tendency. Results are presented as least squares means (LSMEANS) ± the standard error of the mean (SEM). No effect of genotype (hemizygous versus homozygous) on ZsGreen1 abundance was observed (*P* > 0.10).

## Results

### Identification of hemizygous and homozygous CMV-ZsGreen1 transgenic piglets

Presence of ZsGreen1 in the skin of neonatal transgenic piglets was detected visually (Figure 1A) and via immunoblot (Figure 1B), indicative of CMV promoter activity. Even without the UV light, transgenic animals were identifiable via the yellow hue of their skin (data not shown), which has been previously noted in GFP pigs [26]. The genotype of the animals was confirmed by conventional PCR (Figure 1D); a schematic depicting the genotyping method is available in Figure 1C. ZsGreen1 was not detected in the skin of control piglets (Figure 1A & 1B).

**Fig. 1.**
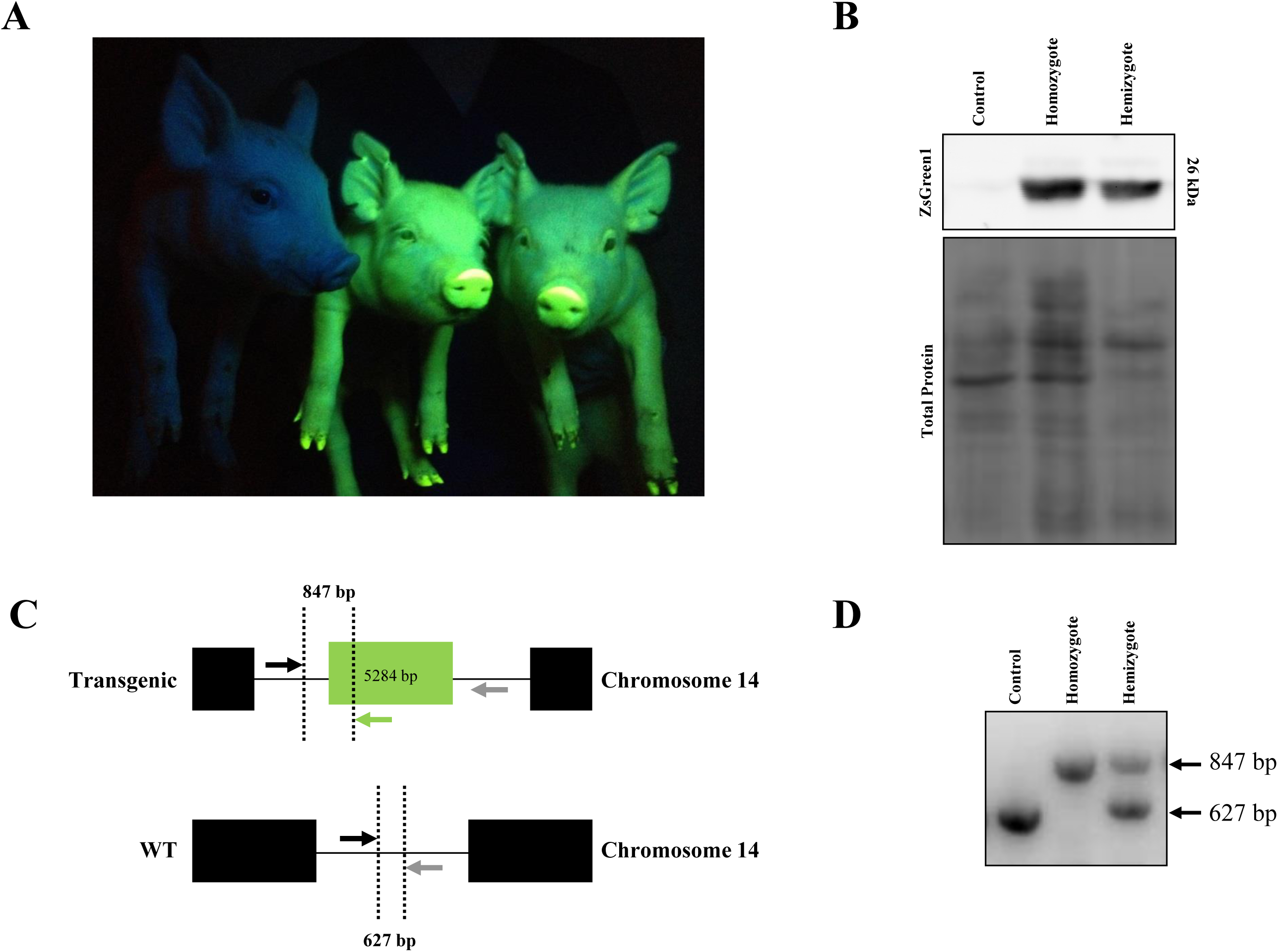
Production of CMV-ZsGreen1 hemizygous and homozygous transgenic pigs. **(A)** Representative photograph of control piglet (left) with homozygous (middle) and hemizygous (right) transgenic siblings. ZsGreen1 was visualized with an ultraviolet light and a Roscolux #15 filter. (**B)** Representative immunoblot of ZsGreen1 in the skin of control piglets (n = 3) as well as hemizygous (n = 3) and homozygous (n = 3) transgenic littermates. Lower panel represents total protein levels within each lane. (**C)** Schematic demonstrating the genotyping method utilized to distinguish between control, homozygous and hemizygous piglets. Detection of monoallelic or biallelic transgene integration was performed by conventional polymerase chain reaction (PCR) using 3 primers. The forward primer, denoted by the black arrow, amplifies a non-coding region upstream of the transgene integration site. A reverse primer (grey arrow) aligns a non-coding region downstream of the transgene and a second reverse primer, indicated in green, detects the transgene. If a transgene is present, the resulting PCR product will be 847 bp; however, in the absence of the transgene, the PCR product will be 627 bp. **(D)** PCR detection of transgene integration and copy number. The presence of only a 847 bp PCR product indicates a homozygous transgenic animal whereas a 627 bp product indicates a control animal. If both 847 and 627 bp products are detected, the genotype is hemizygous transgenic.

### ZsGreen1 fluorescence within tissues of CMV-ZsGreen1 transgenic piglets

ZsGreen1 fluorescence was visually detectable within numerous internal organs of transgenic pigs under UV light; fluorescence was absent in wild-type pig organs (Figure 2A). Fluorescence appeared robust in the intestines (Figure 2A; middle panel), cerebellum (Figure 2B; left panel), anterior pituitary gland (Figure 2B; middle panel) and heart (Figure 2B; right panel), but modest within the stomach (Figure 2A; middle panel). ZsGreen1 was not visually detectable in the cerebrum (Figure 2B; left panel), posterior pituitary gland (Figure 2B; middle panel), liver or spleen (Figure 2A; middle panel). Differential fluorescent patterns were observed within organs as well. In the brain, ZsGreen1 fluorescence was intense within the cerebellum and not visually apparent in the cerebrum (Figure 2B; left panel). In addition, expression of ZsGreen1 appeared greatest in the right ventricle compared with other regions of the heart (Figure 2B; right panel).

**Fig. 2.**
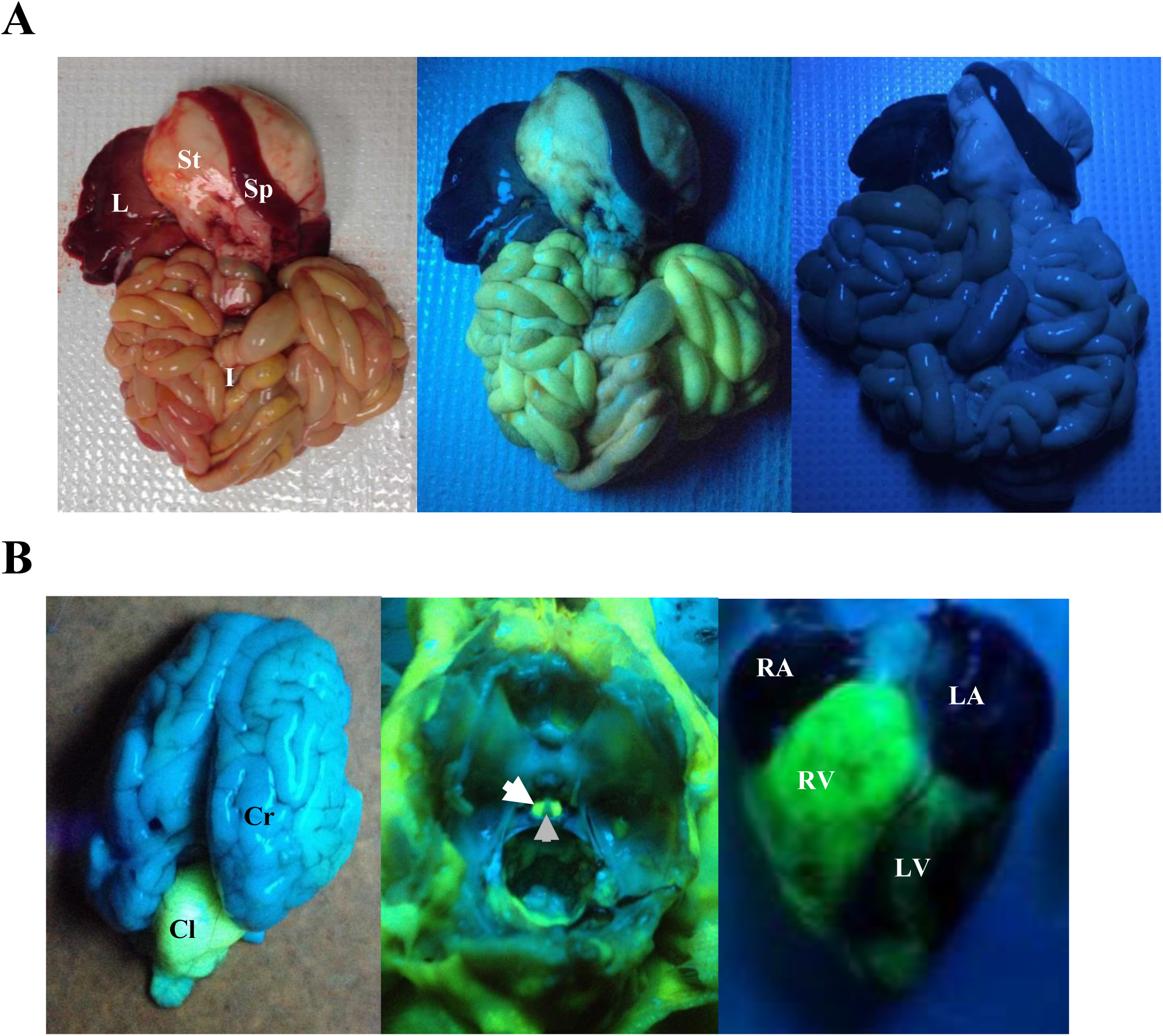
Visualization of ZsGreen1 fluorescence within internal organs of transgenic piglets. **(A)** Representative image of ZsGreen1 fluorescence in the internal organs of control and transgenic pigs. Left panel represents internal organs from a transgenic piglet. Middle panel represents a fluorescent image of internal organs from a transgenic piglet; and right panel is a UV image of wild-type piglet organs. **(B)** Representative fluorescent images of a transgenic piglet brain (left panel), anterior and posterior pituitary glands (within the sella turcica of the skull; middle panel) and heart (right panel). Anterior pituitary gland and posterior pituitary gland are indicated by white and grey arrows, respectively. ZsGreen1 expression was abundant within the right ventricle of the heart (right panel). Abbreviations: L, liver; St, stomach, Sp, spleen; I, intestines; Cr, cerebrum; Cl, cerebellum, RA, right atrium; RV, right ventricle; LA, left atrium; LV, left ventricle. ZsGreen1 fluorescence was detected with UV light and a Roscolux #15 filter.

### Abundance of ZsGreen1 in hemizygous and homozygous porcine organs

#### Neural and endocrine tissues of the brain

There was an effect of tissue type on ZsGreen1 production levels within the brain and nervous system of hemizygous piglets (*P* = 0.0002). Abundance of ZsGreen1 was greatest within the anterior pituitary gland, followed by the cerebellum (*P* < 0.05; Figure 3A). ZsGreen1 levels were lowest in the cerebrum, spinal cord, hypothalamus and posterior pituitary gland (*P* < 0.05), which were not different from each other (*P* > 0.05; Figure 3A). In homozygous piglets, there was also an effect of tissue type on ZsGreen1 abundance (*P* = 0.0396). The anterior pituitary gland expressed more ZsGreen1 than the cerebrum, spinal cord, hypothalamus or posterior pituitary gland (*P* < 0.05; Figure 3B), but was not different from the cerebellum (*P* > 0.05; Figure 3B). ZsGreen1 levels were similar between the cerebellum, cerebrum, spinal cord, hypothalamus and posterior pituitary gland (*P* > 0.05; Figure 3B).

**Fig. 3.**
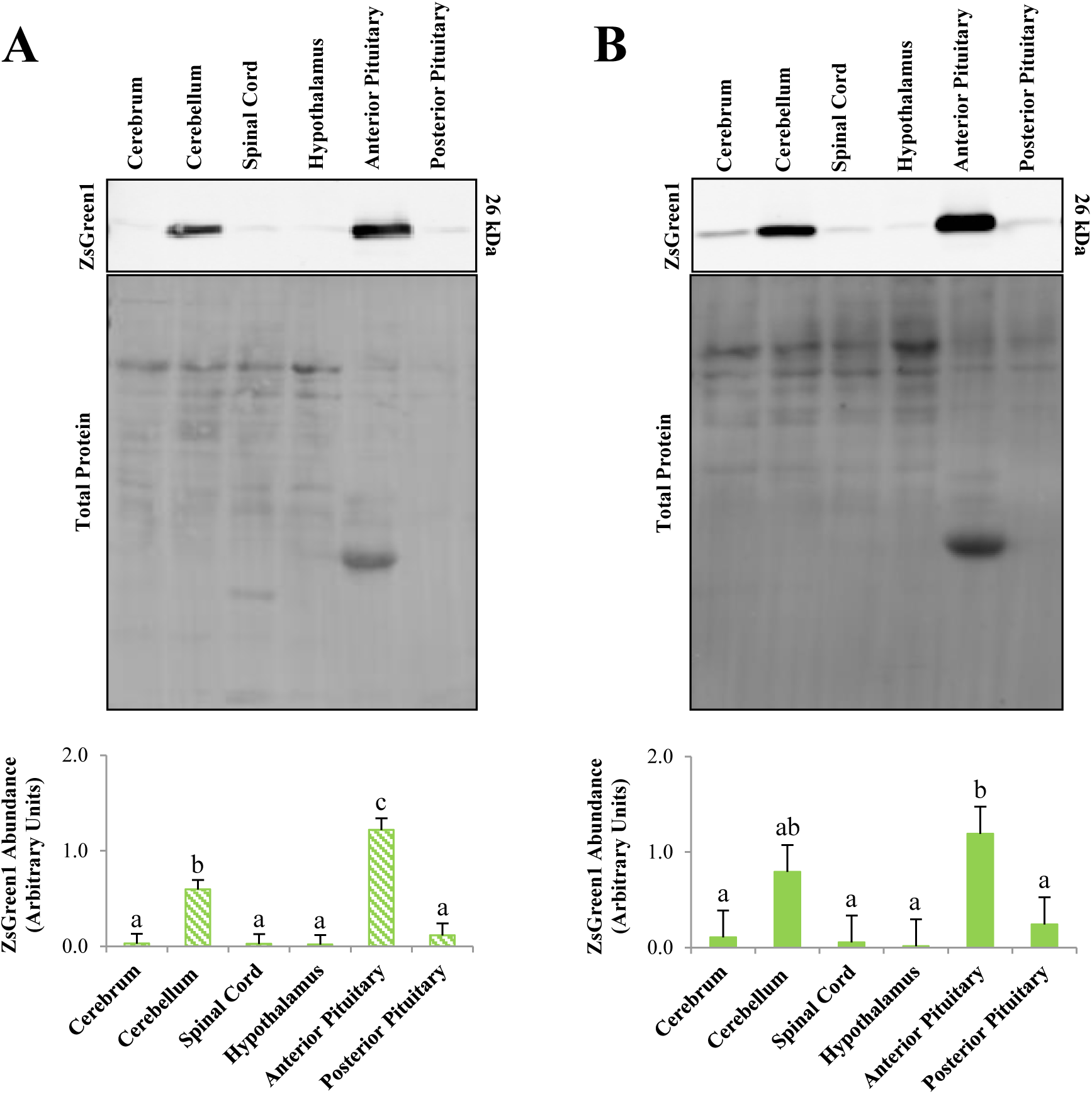
Protein levels of ZsGreen1 within neural and endocrine tissues of CMV-ZsGreen1 pigs. **(A)** Representative immunoblot of neural and endocrine tissue from hemizygous transgenic piglets (n = 3) using an antibody directed against ZsGreen1 (upper panel). Middle panel represents total protein levels that served as the loading control. Quantification of immunoblots revealed differences in relative ZsGreen1 protein levels between tissue types (*P* = 0.0002; lower panel). (**B**) Representative immunoblot of neural and endocrine tissue from homozygous transgenic piglets (n = 3) using an antibody directed against ZsGreen1 (upper panel). Middle panel represents total protein levels that served as the loading control. Quantification of immunoblots revealed a difference in relative ZsGreen1 protein levels between tissue types (*P* = 0.0396; lower panel). ^a,b,c^ Bars with alternate letters differ (*P* < 0.05).

#### Thoracic organs

An effect of tissue type on ZsGreen1 protein abundance was observed among 9 thoracic organs of hemizygous piglets (*P* < 0.0001). The right ventricle produced the most ZsGreen1 in the thoracic region, even when compared with the other chambers of the heart (*P* < 0.05; Figure 4A). ZsGreen1 levels in the right ventricle were greater (*P* < 0.05) than the thyroid, right atrium, left atrium, left ventricle, lung, thymus or lymph node, which were similar (*P* > 0.05; Figure 4A). ZsGreen1 levels within skeletal muscle were intermediate (*P* < 0.05; Figure 4A). In homozygous piglets, there was no effect of tissue type on expression of ZsGreen1 among thoracic organs (*P* = 0.3326; Figure 4B). Although, values for ZsGreen1 abundance were numerically higher in the right ventricle and skeletal muscle compared with other tissues.

**Fig. 4.**
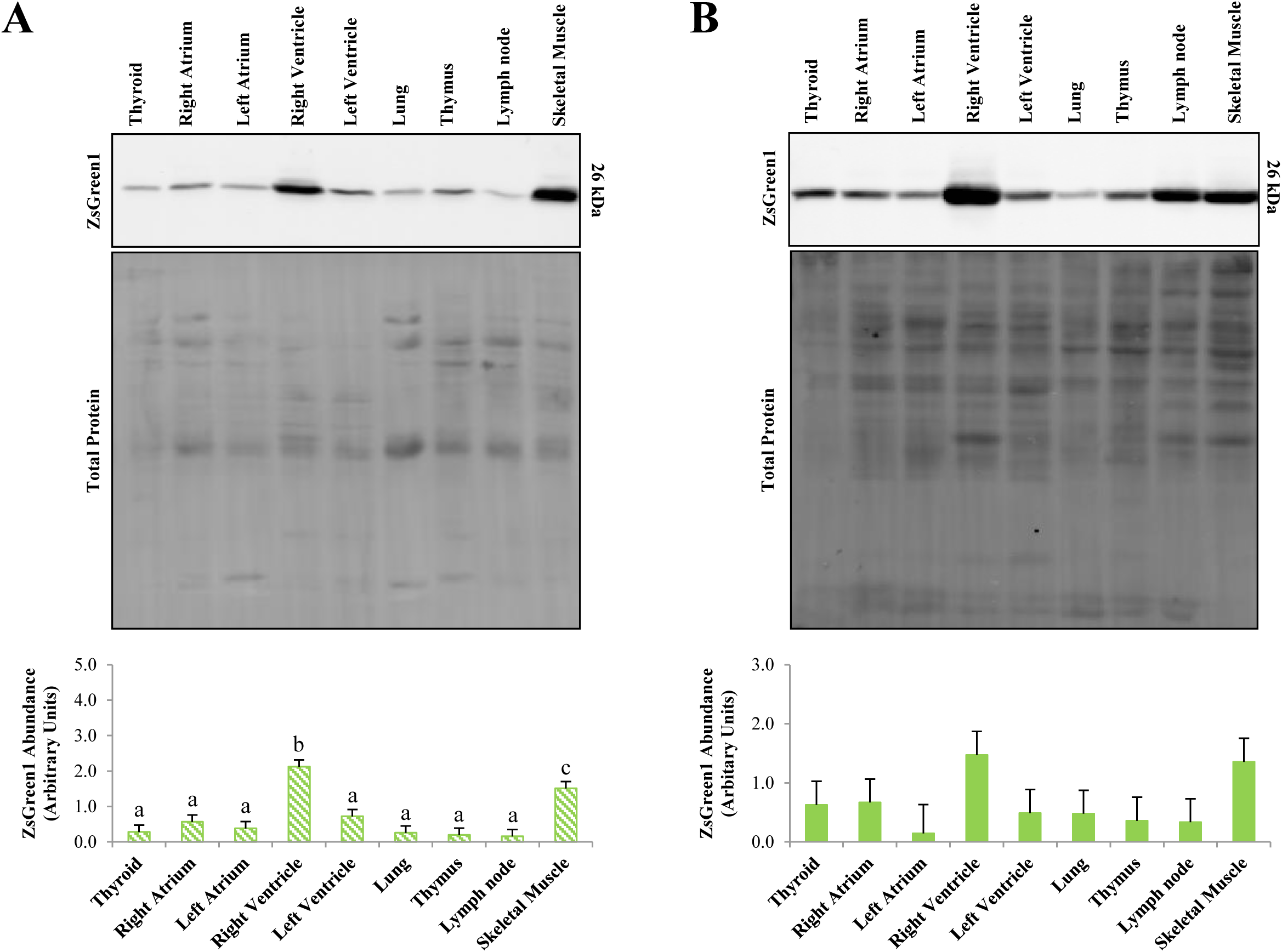
Protein levels of ZsGreen1 within thoracic tissues of CMV-ZsGreen1 pigs. **(A)** Representative immunoblot of thoracic tissue from hemizygous transgenic piglets (n = 3) using an antibody directed against ZsGreen1 (upper panel). Middle panel represents total protein levels that served as the loading control. Quantification of immunoblots revealed differences in relative ZsGreen1 protein levels between tissue types (*P* < 0.0001; lower panel). (**B**) Representative immunoblot of thoracic tissue from homozygous transgenic piglets (n = 3) using an antibody directed against ZsGreen1 (upper panel). Middle panel represents total protein levels that served as the loading control. Quantification of immunoblots indicated that there were no differences in relative ZsGreen1 abundance between tissue types (*P* = 0.3326; lower panel). ^a,b,c^ Bars with alternate letters differ (*P* < 0.05).

#### Digestive Organs

ZsGreen1 protein levels differed across 11 digestive organs of hemizygous piglets (*P* < 0.0001). Levels were greatest in the salivary gland and lowest in the stomach, pancreas, duodenum, jejunum, ileum, colon, spleen, gall bladder, and liver, which were not different (*P* > 0.10). Protein amounts were intermediate in the esophagus (*P* < 0.05; Figure 5A), with the exception of the duodenum which was similar (*P* > 0.05). In homozygous piglets, there was also an effect of tissue type on ZsGreen1 protein abundance (*P* = 0.0212). Expression was greatest in the salivary gland (*P* < 0.05; Figure 5B) compared with the other 10 organs, which were similar (*P* > 0.05; Figure 5B).

**Fig. 5.**
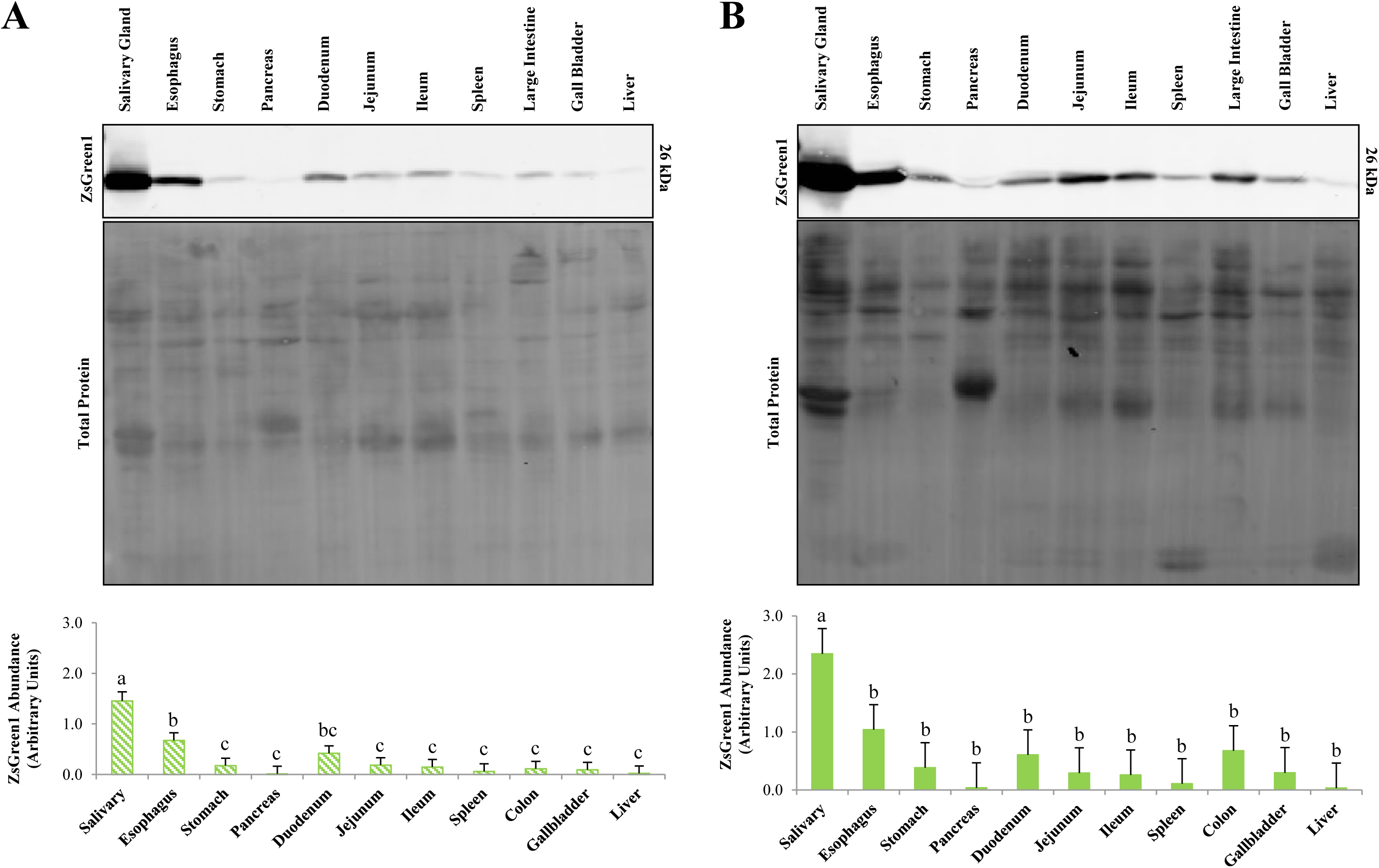
Protein levels of ZsGreen1 within digestive tissues of CMV-ZsGreen1 pigs. **(A)** Representative immunoblot of digestive tissue from hemizygous transgenic piglets (n = 3) using an antibody directed against ZsGreen1 (upper panel). Middle panel represents total protein levels that served as the loading control. Quantification of immunoblots revealed differences in relative ZsGreen1 protein levels between tissue types (*P* < 0.0001; lower panel). (**B**) Representative immunoblot of digestive tissue from homozygous transgenic piglets (n = 3) using an antibody directed against ZsGreen1 (upper panel). Middle panel represents total protein levels that served as the loading control. Quantification of immunoblots revealed differences in relative ZsGreen1 protein levels between tissue types (*P* = 0.0212; lower panel). ^a,b,c^ Bars with alternate letters differ (*P* < 0.05).

#### Renal Organs

An effect of tissue type was observed between 5 renal tissues of hemizygous piglets (*P* = 0.0083). The adrenal medulla produced more ZsGreen1 than the adrenal cortex, kidney medulla, kidney cortex or bladder (*P* < 0.05; Figure 6A), which were similar to each other (*P* > 0.05). In homozygous animals, there tended to be a difference in ZsGreen1 abundance across renal tissues (*P* = 0.0620). Levels were highest in the adrenal medulla and lowest in the kidney medulla, kidney cortex, and bladder (*P* < 0.05; Figure 6B), which were not different from one another (*P* > 0.05). ZsGreen1 levels were similar between the adrenal cortex and all other organs (*P* > 0.05; Figure 6B).

**Fig. 6.**
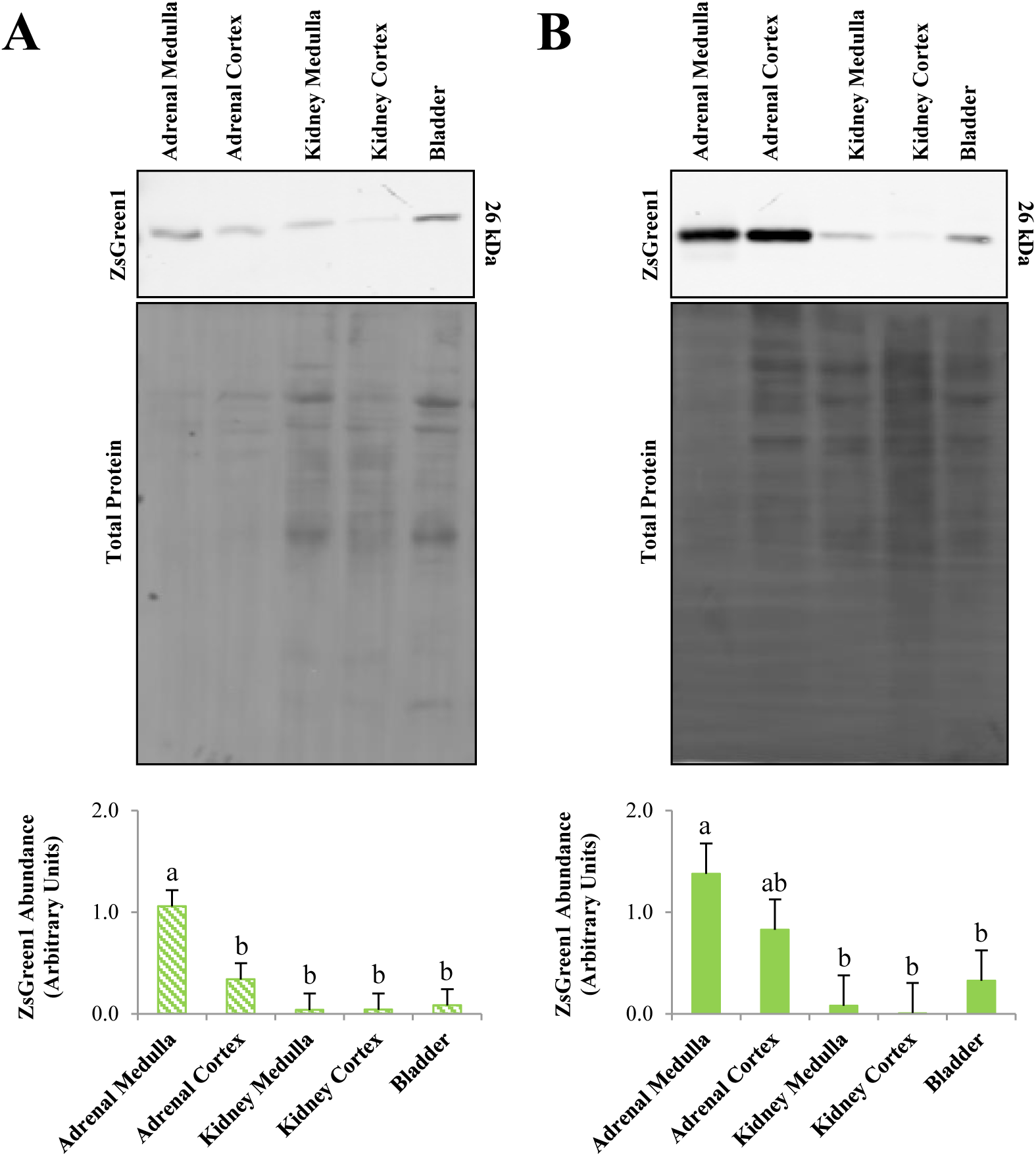
Protein levels of ZsGreen1 within renal tissues of CMV-ZsGreen1 pigs. **(A)** Representative immunoblot of renal tissue from hemizygous transgenic piglets (n = 3) using an antibody directed against ZsGreen1 (upper panel). Middle panel represents total protein levels that served as the loading control. Quantification of immunoblots revealed differences in relative ZsGreen1 protein levels between tissue types (*P* = 0.0083; lower panel). (**B**) Representative immunoblot of renal tissue from homozygous transgenic piglets (n = 3) using an antibody directed against ZsGreen1 (upper panel). Middle panel represents total protein levels that served as the loading control. Quantification of immunoblots revealed a tendency for differences in relative ZsGreen1 protein levels between tissue types (*P* = 0.0620; lower panel). ^a,b,c^ Bars with alternate letters differ (*P* < 0.05).

#### Reproductive organs

No differences in ZsGreen1 expression were detected between the ovary, oviduct, and uterus of hemizygous or homozygous piglets (*P* > 0.05; Figure 7A and 7B).

**Fig. 7.**
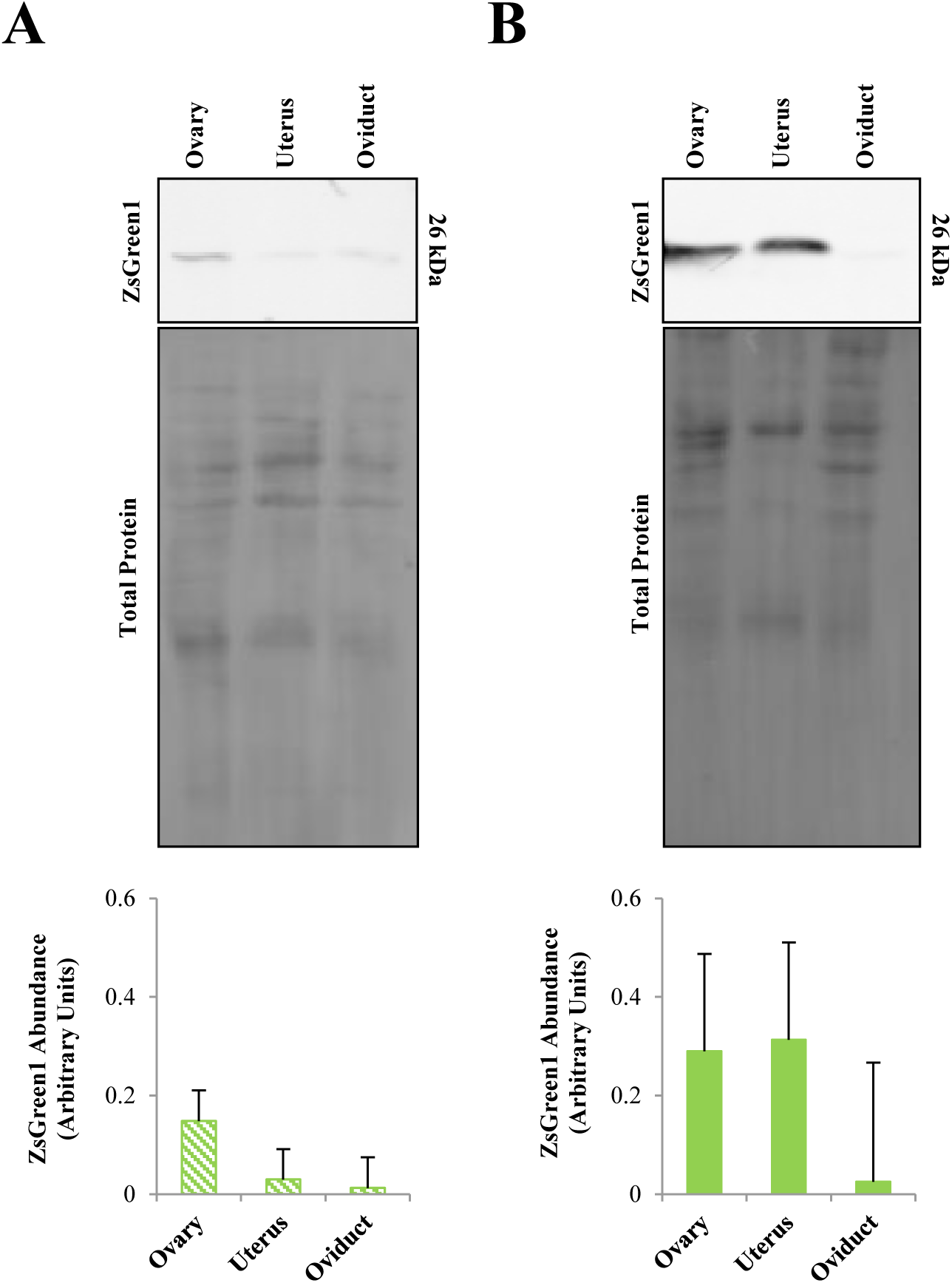
Protein levels of ZsGreen1 within reproductive tissues of CMV-ZsGreen1 pigs. **(A)** Representative immunoblot of reproductive tissue from hemizygous transgenic piglets (n = 3) using an antibody directed against ZsGreen1 (upper panel). Middle panel represents total protein levels that served as the loading control. Quantification of immunoblots indicated that there were no differences in relative ZsGreen1 abundance between tissue types (*P* = 0.3227; lower panel). (**B**) Representative immunoblot of reproductive tissue from homozygous transgenic piglets (n = 3) using an antibody directed against ZsGreen1 (upper panel). Middle panel represents total protein levels that served as the loading control. Quantification of immunoblots indicated that there were no differences in relative ZsGreen1 abundance between tissue types (*P* = 0.4797).

## Discussion

Our study provides a quantitative, spatial analysis of tissue-specific ZsGreen1 protein abundance in swine containing a CMV-ZsGreen1 transgene. In accord with the ubiquitous nature of the CMV promoter, ZsGreen1 protein was detected in every tissue examined via immunoblotting. Consistent with this, another report from our laboratory indicated that exosomes isolated from the milk of hemizygous females are labeled with ZsGreen1 [31]. Our results, however, demonstrate that the level of CMV promoter activity was variable across organs of the neonatal piglet. Others have observed similar results in different animal models expressing a transgene driven by the CMV promoter [14, 25, 26, 32]. Based upon these results, several hypotheses regarding the differential activity of the CMV promoter *in vivo* have arisen. The activity of the CMV promoter within a given cell type may be influenced by integration site [32]; namely, CMV promoter activation may only occur in transcriptionally active regions of the genome [12]. Stochastic silencing can occur when a transgene is integrated near a heterochromatic region, such as a centromere [33], rendering the promoter inaccessible to transcriptional machinery [34]. Stochastic silencing of transgenes can also occur when multiple copies of the transgene integrate in tandem [35], due to local formation of heterochromatin [34]. In support of this hypothesis, Furth et al. [32] observed that two different transgenic mouse lines had marked differences in CMV promoter activity across tissues of the body, despite harboring the same transgene. Likewise, transgenic animals harboring a transgene, but lacking expression of the gene, have been generated [36]. These results could also be due to promoter methylation, which has been shown to silence transgenes *in vitro* and *in vivo* [15, 16, 21, 37]. Conversely, Vasey et al. (2009) hypothesized that integration site does not readily affect CMV promoter activity because a similar activity profile was detected in pigs and chickens harboring the same transgene, albeit integrated in different chromosomal regions [26, 27]. The activity patterns were also similar in multiple transgenic chicken lines expressing different reporter genes directed by the CMV promoter [25]. Our data support these results as a similar expression pattern was observed in CMV-ZsGreen1 pigs, despite a different integration site and transgene.

In addition to chromatin status, the repertoire of transcription factors and co-regulators present within the cell dictate CMV promoter activity [12]. Mella-Alvarado et al. (2013) hypothesized that CMV promoter activity is robust in the organs that CMV infects most voraciously (brain, eye, spinal cord, pancreas, kidney, testis, ovary, skin, cartilage, and skeletal muscle), reflective of optimal transcription factor expression. Moreover, Furth et al. [32] suggested that certain transcription factors may only be able to interact with euchromatin, whereas others retain functionality with heterochromatin [32]. In exocrine glands, Vasey et al. [26] proposed that the transcription factor, nuclear factor kappa-light-chain-enhancer of activated B cells (NF-kB), is a major driver of CMV promoter activity. NF-kB is highly expressed in exocrine cells, classically regulates the expression of proinflammatory genes [38], activates in response to cellular insult such as viral infection [38], and is a potent regulator of the CMV promoter [12, 39-41]. In addition to NF-kB, numerous transcription factors have been implicated in the regulation of the CMV promoter, including ying yang 1 [12], activator protein 1 and its major subunits (c-Fos and c-Jun) [41], p53, cAMP response element binding nuclear transcription factor/activating transcription factor, retinoic acid receptor [42], specificity protein 1 [11], nuclear factor I, serum response factor, Elk-1 and CCAAT/enhancer-binding protein [13]. Thus, it is apparent that the CMV promoter has adapted to respond to a variety of cell signals in order to allow for the promiscuous infection of host cells [12].

Consistent with elevated activity of the CMV promoter in exocrine glands, ZsGreen1 protein levels were increased in the salivary glands of hemizygous and homozygous transgenic piglets, as previously noted in CMV-GFP pigs [26]. Enhanced CMV promoter activity was reported in other exocrine tissues of the pig, including the glandular epithelia of the snout, sebaceous glands of the skin and Harderian gland of the eye [26]. In the exocrine pancreas, however, we did not detect elevated ZsGreen1 levels in either hemizygous or homozygous transgenic piglets, conflicting with other reports of increased CMV promoter activity within the porcine pancreas [26, 27]. This discrepancy could be related to age-specific activity levels of the CMV promoter, as noted in zebrafish [14]. In the current study, neonatal (1 d old) piglets were utilized, whereas others examined CMV promoter activity in the pancreases of older (2 mo to 1 yr) pigs [26, 27]. Likewise, our laboratory detected robust ZsGreen1 abundance in the pancreas of 1 mo old piglets (unpublished data). Interestingly, the exocrine pancreas is largely quiescent in the neonatal piglet [43-47]; production of pancreatic juices and enzymes increase dramatically after weaning [43-45]. In the liver, another exocrine organ that secretes bile made by hepatocytes [48], we also detected little CMV promoter activity. Others have reported similar levels of CMV promoter activity in porcine hepatocytes [26] and chicken liver [25]. In contrast, GFP was robustly driven by the CMV promoter in hepatocytes of zebrafish [14]. Thus, it appears that the CMV promoter is not uniformly effective in all exocrine organs and can vary in activity across species.

In the current study, abundance of ZsGreen1 was also compared between homozygous and hemizygous animals. Based on our visual observation of ZsGreen1 production in the skin of hemizygous and homozygous piglets (Figure 1A), we hypothesized that an additional transgene would yield more ZsGreen1 protein in the organs of homozygous animals. However, no genotype effect was detected based upon quantification of immunoblots for ZsGreen1. These results may be due to monoallelic expression, which frequently occurs in autosomes [49] and has also been described in other transgenic animal models [34, 50]. Conversely, translation of ZsGreen1 transcripts may be impaired in homozygous animals. For example, post-transcriptional silencing of high expressing transgenes is common in homozygous, but not hemizygous, plants [51-54] and appears to be mediated by short interfering RNA [51]. Many have hypothesized that doubling transcription of a transgene yields copious mRNA transcripts, which breaches an unknown threshold that triggers cellular processes to suppress transgene translation [52, 55-58]. Regardless of the mechanism, transgene copy number did not significantly affect production of ZsGreen1 protein in this study.

To our knowledge, these animals represent the first pig model to express ZsGreen1 (Clontech, Palo Alto, CA). This protein, originally discovered by Matz et al. [59], is derived from Anthozoa coral reef (*Zoanthus* species). The variant, ZsGreen1, is a modification of the original protein (zFP506) to maximize expression, solubility and prompt chromophore maturation (Clontech, personal communication). Interestingly, it is both structurally and functionally distinct from other fluorescent proteins, like GFP [59, 60]. Although both fluorescent proteins are configured in a β-barrel structure [59], ZsGreen1 exists as a tetramer, whereas GFP (derived from jellyfish) is a monomer that frequently dimerizes for solubility [61]. Interestingly, ZsGreen1 is only 26% homologous to GFP and each ZsGreen1 monomer is 231 amino acids in length with a predicted molecular weight of approximately 26 kDa [59]. Functionally, ZsGreen1 displays extremely bright fluorescence and high signal-to-noise ratio; in fact, yeast cells overexpressing ZsGreen exhibited 8.6-fold more fluorescence than those expressing GFP [60]. In another study, ZsGreen1 had the greatest relative fluorescence compared with 5 other fluorescent proteins, including GFP [59].

Notably, this ZsGreen1 swine model could serve as a useful tool for genome editing applications. This model is well suited for re-engineering the transgene locus since expression of the transgene is readily detectable, the transgene is present in a single copy and the integration site has been identified [4]. For example, this swine line could be used for testing somatic cell gene editing using CRISPR/Cas9 systems or base editor nucleases by targeting the ZsGreen1 transgene sequence. In addition, the model can be a useful tool for testing germline genome editing approaches *in vivo* such as Genome editing via Oviductal Nucleic Acids Delivery (GONAD), initially established in the mouse [62]. Finally, this pig model can be used for re-engineering the transgenic CMV-ZsGreen1 locus to make it suitable for other applications. For example, the ZsGreen1 coding sequence can be disrupted to create a frameshift mutation in ZsGreen1, creating a model to serve as a reporter for homology directed repair genome editing studies. Alternatively, a floxed red fluorescent protein could be inserted upstream of the ZsGreen1 cassette to serve as a Cre recombinase reporter, similar to what is popularly used in mouse Cre-LoxP studies [63]. Re-engineering of the locus can be achieved via Easi-CRISPR, a highly efficient genome engineering method recently described in mice [64, 65].

## Conclusions

This study demonstrates that the CMV promoter is ubiquitously active in all porcine tissues types examined; however, the level of activity is tissue-specific. The CMV promoter potently drives transgene expression in many exocrine (e.g., salivary gland) and non-exocrine organs (e.g., skin, anterior pituitary gland and muscle) of the neonatal pig. In certain organs (e.g., liver, kidney), however, activity of the CMV promoter is less robust. Ultimately, this report elucidates the tissue-specific activity of the human CMV promoter in swine and provides critical information for researchers seeking an effective promoter to drive transgene expression in the neonatal pig. Moreover, these animals represent a novel resource for scientists in need of ZsGreen1-labeled porcine organs. Lastly, this CMV-ZsGreen1 swine model will also serve as a valuable tool to advance genome editing research.

## Abbreviations

CMV: cytomegalovirus promoter
DNA: deoxyribonucleic acid
GFP: green fluorescent protein
NF-kB: nuclear factor kappa-light-chain-enhancer of activated B cells
LSMEANS: least squares means
PCR: polymerase chain reaction
SDS: sodium dodecyl sulfate
SEM: standard error of the mean
UV: ultraviolet

## Declarations

### Acknowledgements

The authors wish to thank Ginger Mills for pig husbandry as well as Scott Kurz, Guilherme Cezar, Kyle Regan and Amanda Lambrecht for technical assistance. The authors would also like to express gratitude to Lelanya Yates for aiding in the dissections.

## Availability of data and materials

The datasets used and/or analyzed during the current study are available from the corresponding author on reasonable request.

## Authors’ contributions

AD, RC and BW conceived of the study and designed the experiment. AD, RC and EP collected the data. AD analyzed the data. AD, RC, GC and BW interpreted the data. AD and RC drafted the manuscript. All authors read, revised and approved the final manuscript.

## Ethics approval and consent to participate

All animal procedures were conducted in accordance with the University of Nebraska-Lincoln Institutional Animal Care and Use Committee.

## Consent for publication

Not applicable.

